# A Repetitive Modular Oscillation Underlies Human Brain Electric Activity

**DOI:** 10.1101/072538

**Authors:** Arturo Tozzi, James F. Peters, Norbert Jaušovec

## Abstract

The modular function j, central in the assessment of abstract mathematical problems, describes elliptic, intertwined trajectories that move in the planes of both real and complex numbers. Recent clues suggest that the j-function might display a physical counterpart, equipped with a quantifiable real component and a hidden imaginary one, currently undetectable by our senses and instruments. Here we evaluate whether the real part of the modular function can be spotted in the electric activity of the human brain. We assessed EEGs from five healthy males, eyes-closed and resting state, and superimposed the electric traces with the bidimensional curves predicted by the j-function. We found that the two trajectories matched in more than 85% of cases, independent from the subtending electric rhythm and the electrode location. Therefore, the real part of the j-function’s peculiar wave is ubiquitously endowed all over normal EEGs paths. We discuss the implications of such correlation in neuroscience and neurology, highlighting how the j-function might stand for the one of the basic oscillations of the brain, and how the still unexplored imaginary part might underlie several physiological and pathological nervous features.

**SIGNIFICANCE STATEMENT:** Our results point towards the brain as ubiquitously equipped with j-function’s oscillations, which movements take place on the plane of the complex numbers. It means that there must be, in brain electric activity, also a veiled complex part, which can be assessed with the help of imaginary numbers. The modular j-function provides further dimensions to the real numbers, in order to enlarge their predictive powers: it suggests the possible presence of hidden (functional or spatial) brain extra-dimensions. Furthermore, j-oscillations could be disrupted during pathologies, paving the way to novel approaches to central nervous system’s diseases.

## INTRODUCTION

The j-function describes intertwined, repetitive elliptic trajectories taking place on the so-called *plane of the complex numbers* (McKeague, 2011). Such plane can be exemplified as a plot displaying on the x axis the real numbers (e.g., 1, 2, −1, −2) and on the y axis the imaginary part (e.g., a component that satisfies the equation *i*^2^ = −1). The intersection of a real and an imaginary number in the graph is termed a *complex number* (**Figure 1A**). The role of complex numbers in mathematics is to enlarge the concept of the one-dimensional number line to the two-dimensional complex plane, allowing the description of real numbers with an additional coordinate. This trick is very useful not just in mathematics, but also in the other disciplines, from fluid dynamics to electrical engineering and computer graphics, from signal analysis to quantum mechanics, and so on (Rankin, 1977; Apostol, 1990).

**Figure 1.**
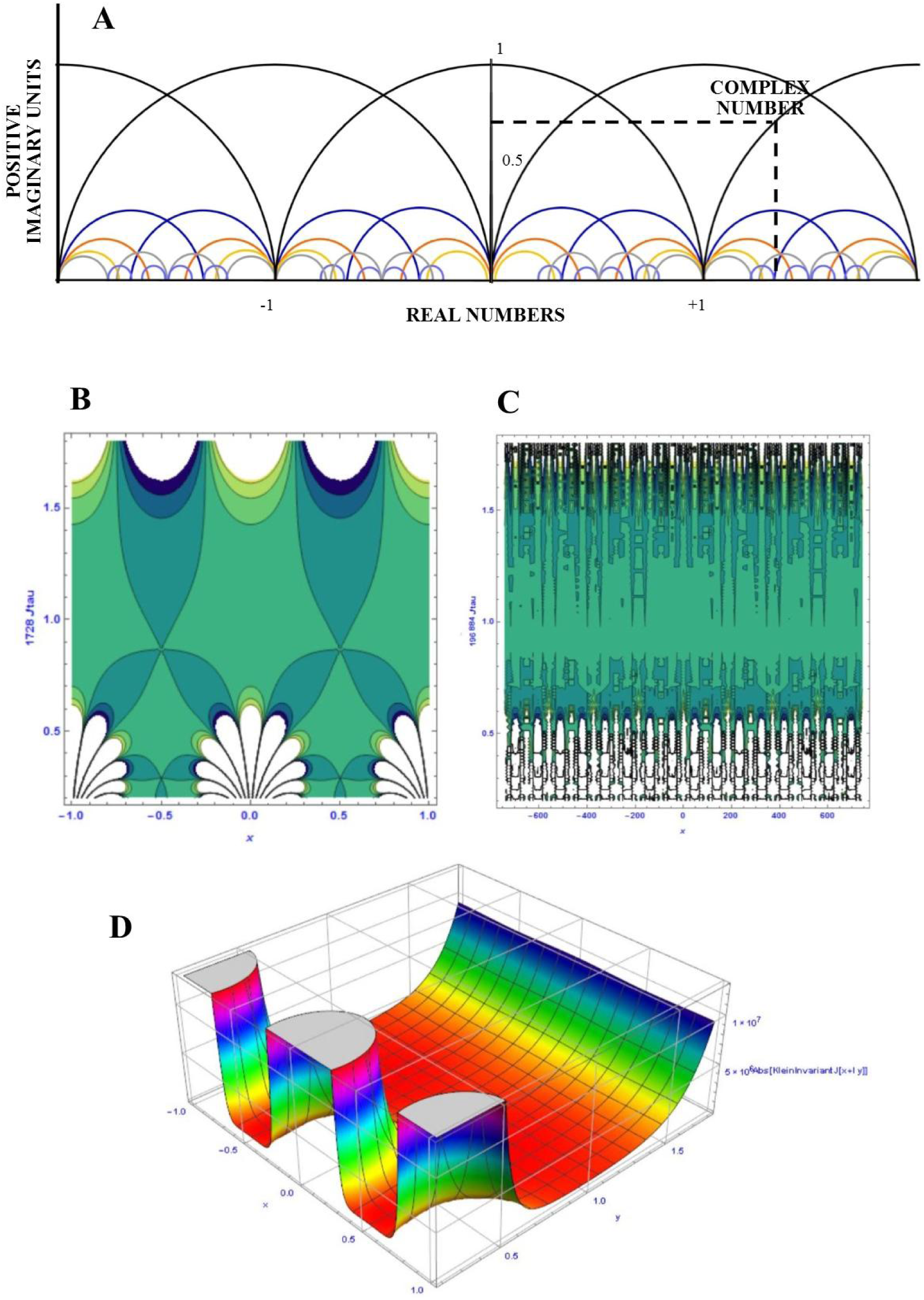
Plots of Klein j-functions and their different possible paths. **Figure 1A** depicts the general scheme of the elliptic oscillations on the upper half plane of the complex numbers. For sake of clarity, the locations of imaginary units and of real and complex numbers are provided. **Figure 1B** and **1C** show the 2D plots achieved for j=1728 and 196,884, respectively. Note the different coarse-grained appearance of the two plots. **Figure 1D** displays the 3D plot, in case of j=196,844. The plots in **Figures 1B, 1C** and **1D** were obtained using Mathematica^®^ 10.

The j-function is plotted on the upper half of the complex plane, e.g., it can be drawn just starting from positive imaginary numbers. The pattern depicted by the j-function is called modular, e.g. in simple terms, it looks like curved trajectories intersecting one each other in a recurrent and fractal pattern. The j-function exhibits the peculiarity that, contrary to other modular functions that can be mapped in all their points on planes, is equipped with cusps, e.g., single points where a trajectory starts to move backward.

It has been speculated that modular functions, and in particular the j-function, might have physical counterparts in our Universe, because they seem to be indirectly linked with string theories that describe the microscopic features of cosmos (Gannon, 2006; Duncan and Frenkel, 2011; Eguchi et al., 2011; Duncan et al., 2015). In particular, it has been suggested that the j-function might count the ways strings can oscillate at each one of their energy levels. Therefore, modular functions might be more widespread than thought and could be an underlying, hidden rhythm that dictates the oscillatory activities of different physical and biological systems. In this paper we evaluated the possibility that brain electric rhythms encompass the j-function. Indeed, j-function and brain activity display, at first sight, many common features: they are both modular, bot sinusoidal, both symmetric (Tozzi and Peters 2016a) and both can be described in terms of Fourier transform. In such a vein, we assessed EEG patterns collected from adult controls during eyes closed resting state, looking for the j-function’s features. We demonstrated that the j-function is endowed in the very structure of human brain electric spikes, giving rise, at least in normal subjects, to reiterating oscillations equipped with a highly predictable pattern. Furthermore, we discuss the implications, both in basic neuroscience and in medical science, of our finding.

## METHODS

### Subjects

The sample included 5 right-handed male students, undergraduates (age =21 years). They were recruited from a large pool of individuals who participated in different experiments conducted at Department of Psychology, University of Maribor lab, in the last 2 years (e.g., Tement, et al., 2016). The participants were tested with the WAIS-R test battery (Wechsler, 1981; M = 100.60; SD = 1.14). The experiment was undertaken with the understanding and written consent of each subject.

### Procedure, EEG recording and quantification

The participants were seated in a reclining chair while their resting eyes closed EEG was recorded for 5 min. Afterwards they solved several tasks which were part of different studies described above. A Quick–Cap with sintered (Silver/Silver Chloride; 8 mm diameter) electrodes was fitted to each participant. Using the Ten-twenty Electrode Placement System of the International Federation, the EEG activity was monitored over nineteen scalp locations (Fp1, Fp2,F3, F4, F7, F8, T3, T4, T5, T6, C3, C4, P3, P4, O1, O2, Fz, Cz and Pz). All leads were referenced to linked mastoids (A1 and A2), and a ground electrode was applied to the forehead. Vertical eye movements were recorded with electrodes placed above and below the left eye. Electrode impedance was maintained below 5 kW. EEG was recorded using NeuroScan Acquire Software and a SynAmps RTamplifier (Compumedics, Melbourne Australia), which had a band-pass of 0.15–100.0 Hz. At cutoff frequencies the voltage gain was approximately –6 dB. The 19 EEG traces were digitized online at 1000 Hz with a gain of 10x (accuracy of 29.80 mV/LSB in a 24 bit A to D conversion) and stored on a hard disk. Offline EEG analysis was performed using NeuroScan Scan 4.5 software (Compumedics, Melbourne, Australia). IAF for each subject was determined based on their resting EEG (Angelakis, et al., 2004). First, power spectra were calculated over the entire epoch length of 11 s (11,132 data points) and automatically screened for artifacts. Second, an automatic peak detection procedure found the highest peak (maximum alpha power) in a 7 to 14 Hz window (frequency steps of 0.09 Hz) separately for each lead. This method was used because it is more adequate for studying endophenotypic qualities during resting (eyes closed) EEG sessions (Klimesch et al., 1993; Hooper, 2005) than the peak frequency center of gravity method proposed by Klimesch et al. (1993), which is used for eyes open conditions. Third, IAF was computed for each channel and averaged over anterior (Fp1, Fp2, F3, F4, F7, F8, Fz), and posterior (T5, T6, P3, P4, O1, O2, Pz) locations ( M = 9.88; SD = 0.34).

For our study, we examined artifact-free sequences of 5 seconds for every eye closed resting individual, from three scalp locations: CZ-, FZ- and PZ-. The EEG traces were filtered for different frequencies, in order to elucidate whether the j-function could be endowed in more than one brain rhythm. IAF between subjects did not differ significantly, therefore the resting eyes-closed EEG recordings were filtered in 5 fixed frequency bands: delta [1.0 – 3.9 Hz]; theta [4.0 – 7.9 Hz]; alpha [8.0 – 12.0 Hz]; beta [15.0 – 31.0 Hz] and gamma [32.0 – 49.0 Hz]. We used a band-pass Infinite Impulse Response (IIR) filter with a 48 dB/octave rolloff.

### Looking for j-function in EEGs

The modular function j is the generator or hauptmodul of the genus zero function field (**Figure 1**). Originally, the j-function was defined by (Lehmer, 1942) by

*j*(*τ*) 1728*J*(*τ*), whose Fourier development is
*j*(*τ*) = *e*^−2*πir*^ + 744 + 196884*e*^2*πir*^ + 21493760^4*πir*^ +…, where
I[τ]> 0 is the half-period ratio and

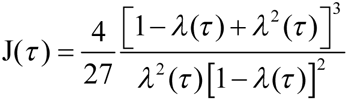

is Klein’s absolute invariant and λ(τ) is the eliptic lambda function (Piezas)

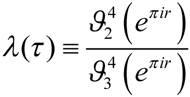, where
ϑ_*i*_(0, *q*) are Jacobi theta functions and
*q* = is the nome and 1728 = 12^3^.

In case of a j-function endowed in brain electric spikes, we would be able to detect just the waves depicting the real part of the modular function, because the imaginary part lies in a complex dimension (either numerical, or functional) that is undetectable by our senses and our technological tools, unless you explicitly look for it. In particular, our goal was to assess whether the EEGs’ bidimensional lines match with the bidimensional real part of the j-function. At first, we built a bidimensional pattern of the j-function. The pattern (**Figure 2A**) describes a curve conventionally moving from T0 to T1, from left to right. The most important parameter to take into account for our purposes is the distance between two peaks, that, being recursive, is the best testable feature of two-dimensional j-curves. The frequency and the amplitude of the sinusoidal oscillations are less important, because, being the modular function fractal, it can be detected at different spatial and temporal coarse-grained scales. Likewise, because the EEG waves are equipped with multiple, unpredictable deformation arising by multiple intertwined interactions, the j-pattern could be nested in different amplitudes and frequencies.

**Figure 2.**
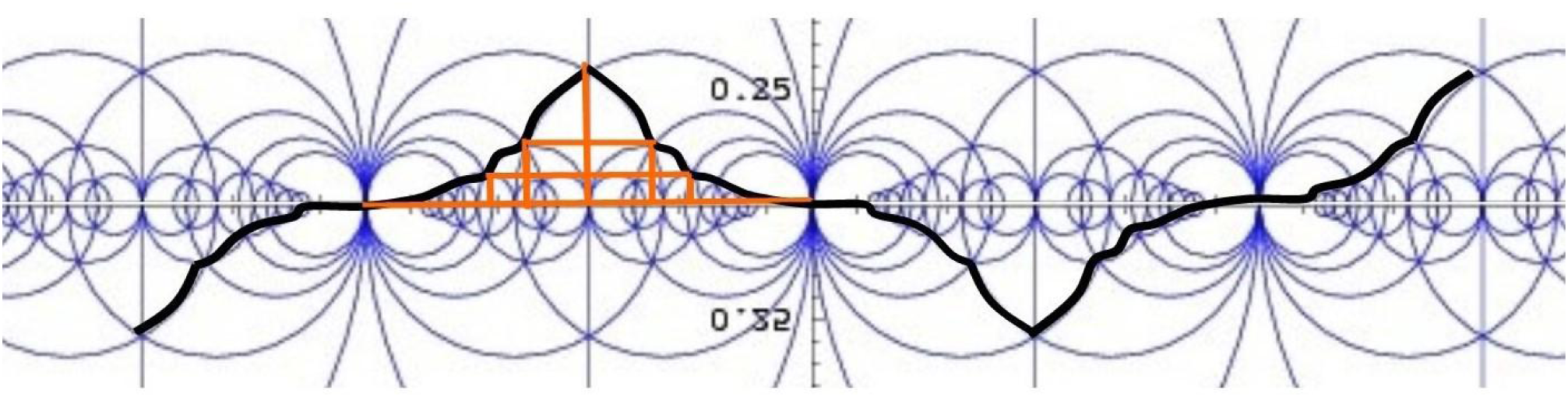
Bidimensional pattern of a j-function, displaying the real part of j-oscillations. The black line depicts the j-pattern correlated with EEGs traces.

We correlated the predicted j-pattern with the above mentioned EEG traces from adult controls. In particular, we evaluated the percentage of the two oscillations’ superimposition. We used a t-student test for statistical analysis.

## RESULTS

The predicted j-pattern, standing for the real part of the j-function, matched the EEG plots recorded from all the normal healthy, resting, eyes-closed males (**Figure 3**). In particular, the modular j-function is correlated with the distance among the peaks. J-function and EEG traces’ superimposition was found in all the subjects and derivations. Taking into account all the frequencies and all the derivations, EEGs oscillations match the pattern of the predicted j-function in a high number of plots (Mean% ± SD: 87.6 ± 5.54). Although not significant, the slower waves (delta, theta and alpha) exhibited a slightly higher correlation (93% vs. 81%) than the faster ones (beta and gamma). It could depend on a more marked intertwining of oscillations in faster frequencies, that partially interferes with the detection of j-function’s typical patterns. No statistical differences were found among different electrodes localizations. In sum, we found that the real part of the j-function’s peculiar wave is ubiquitously endowed all over normal EEGs traces, independent of electric rhythms and electrode localizations.

**Figure 3.**
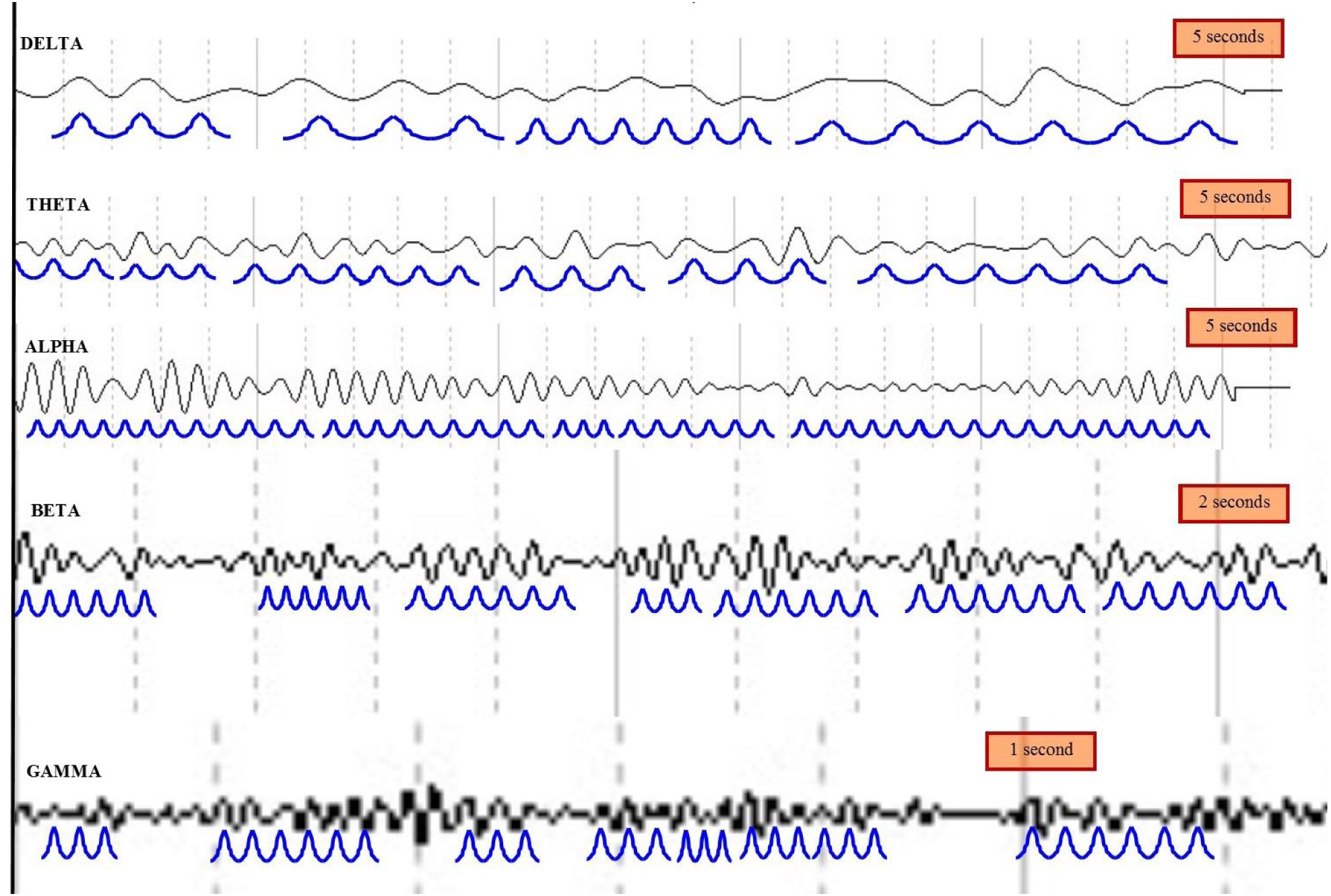
Relationships between the j-function pattern (blue waves) and normal EEG rhythms (black waves) in the FZ-derivation of a single male control. The predicted blue curves match with the normal EEG oscillations almost everywhere. Delta, theta and alpha traces have the same resolution for the y (microV) axis, which gives a rather unclear view for the beta and gamma waves, usually in the < 1mV range. Therefore, beta and gamma frequencies are depicted in Figure at higher magnification and with a different temporal scale.

## CONCLUSIONS

We demonstrated that normal EEG’s waves encompass the real part of a modular function. Our results point towards the human brain as equipped with a pattern resulting from elliptic, intertwined oscillations shaped exactly in guise of the waves described by the j-function. Such oscillations endowed in the brain spikes result from the delicate interaction of the countless moving trajectories that are at the very heart of modular functions. The presence in EEGs of the j-function, which movements take place on the so-called plane of the complex numbers, means that there must be, in brain electric activity, also a still unexplored complex part. While current neuro-techniques are able to detect and describe just the real part of nervous function, e.g., the part quantifiable through real numbers, the complex part is hidden from our observation, because it can be described just with the help of an imaginary component. It means that electric features linked with elusive brain functions, such as, e.g., long-distance interactions, consciousness and so on, could be unnoticed in the EEG traces. They could be assessable, if we add further parameters able to assess neural data sets not just in terms of real numbers, but also of complex ones. In such a vein, it must be taken into account that EEG techniques are able to assess just the part of the j-function that lies very close to the x axis, therefore missing the extensive part of the upper half plane of complex functions that lies farther away from the axis of real numbers. Because the j-function is a mathematical tool which provides further dimensions to the real numbers in order to enlarge their predictive powers, it could be hypothesized that the mathematical relationship and superimposition between modular functions and EEG spikes might be linked with the presence of further brain dimensions, undetectable by our 3D analyses. Indeed, the possibility of hidden functional or spatial nervous dimensions has been recently discussed (Tozzi and Peters, 2016b). The Authors described the occurrence, at least during spontaneous brain activity, of multidimensional toruses, where trajectories take place in guise of particles traveling on donut-like manifolds.

Although the possible presence of nested (real and the imaginary) parts of the j-function has not yet been explored in physical and biological phenomena, there are good reasons for hypothesizing that such j-oscillations could be widespread. For example, the Monstrous Moonshine conjecture suggests a puzzling relationship between one of the terms in the Fourier expansion of j(τ), which value is 19884, and the simple sums of dimensions of irreducible representation of the Monster group M, which is 196883 (Conway and Norton, 1979). On the other hand, the Monster group has been correlated with physical features underlying our Universe (Frenkel et al., 1984; Duncan and Frenkel, 2011). Furthermore, the j-function displays both (spatial) fractal-like structure, and (temporal) power laws, in touch with the description of many physical and biological systems (including the brain) in terms of nonlinear systems equipped with scale-free behavior (Linkenkaer-Hansen, et al., 2001; Newman, 2005; Milstein et al., 2009; He et al., 2010; Tozzi, 2015). To give other examples, it has been recently proposed that symmetry breaks occur during brain function (Tozzi and Peters 2016a) and that the nervous activity might be described in terms of a gauge theory (Sengupta et al., 2016). A gauge theory states that systems’ general symmetries can be broken by local perturbations and then restored by the superimposition of gauge fields. The modular function j, with its repetitive and highly conserved pattern, is a very good candidate for the role of a general symmetry of the brain (and other systems).

Once established that the j-function’s is ubiquitously endowed all over normal EEGs traces during resting state, eyes open conditions, the further steps could be: a) to evaluate task related EEGs, and b) to superimpose the j-pattern to nervous oscillations other than electric spikes, such as the ones detected by, e.g., fMRI or PET. Once elucidated the role of j-function in physiological brain oscillations, it will be worthwhile to look for possible j-function’s alterations in pathological states. Indeed, it could be hypothesized that the recurring j-oscillations are disrupted during pathologies, paving the way to novel approaches to brain diseases.

